# Comparison of SARS-CoV-2 VOC 202012/01 (UK variant) and D614G variant transmission by different routes in Syrian hamsters

**DOI:** 10.1101/2021.03.26.437153

**Authors:** Sreelekshmy Mohandas, Pragya D Yadav, Dimpal Nyayanit, Anita Shete-Aich, Prasad Sarkale, Supriya Hundekar, Sanjay Kumar, Kavita Lole

**Affiliations:** Indian Council of Medical Research-National Institute of Virology, Pune, Maharashtra, India, Pin-411021; Department of Neurosurgery, Command Hospital (Southern Command), Armed Forces Medical College, Pune, Maharashtra, India, Pin-411040

**Keywords:** SARS-CoV-2, VOC 202012/01, UK variant, D614G variant, transmission, Syrian hamsters

## Abstract

Many SARS-CoV-2 variants of concern has been reported recently which were linked to increased transmission. In our earlier study on virus shedding using VOC 202012/01(UK variant) and D614G variant in hamster model, we observed significantly higher viral RNA shedding through nasal wash in case of UK variant. Hence, we compared the transmission of both the UK and D614G variant by various routes in Syrian hamsters to understand whether the high viral RNA shedding could enhance the transmission efficiency of the variant. The current study demonstrated comparable transmission efficiency of both UK and D614G variants of SARS-CoV-2 in Syrian hamsters.

## Main Text

SARS-CoV-2 virus has accumulated numerous genetic changes on circulation all over the world. Some of these mutations have the potential to change the virus characteristics like infectiousness, transmissibility, severity of disease and can have impact on diagnostics, vaccines and therapeutics ^1^. SARSCoV-2 variant with a D614G substitution which emerged in the first quarter of the year 2020 supplanted the initial SARS-CoV-2 strain showing its increased fitness to become the dominant strain circulating globally. Recently, many variants of concern (VOC) were reported from United Kingdom, South Africa, Brazil etc. which were linked to increased transmission, disease severity and vaccine escape mutants ^2^. To categorize a variant as VOC, various risk elements like increased transmissibility, morbidity, mortality, immunity escape factors needs to be studied. Scanty information is available on these aspects about the variants.

In our earlier study on virus shedding using VOC 202012/01(UK variant) and D614G variant in hamster model, we observed significantly higher viral RNA shedding through nasal wash in case of UK variant ^3^. Direct contact, aerosol and fomite routes of transmission of SARS-CoV-2 has been established in hamster model ^4^. Here we have compared the transmission of both the UK and D614G variant by various routes in Syrian hamsters to understand whether the high viral RNA shedding through nasal cavity in hamsters infected with UK variant could enhance the transmission efficiency of the variant.

The study was approved by the Institutional Animal Ethics and Biosafety Committee of Indian Council of Medical Research (ICMR) -National Institute of Virology (NIV), Pune and all the experiments were performed as per the institute guidelines. The study was performed in the containment facility with a total of 36 male hamsters of 6-8 week age housed in individually ventilated cages. Nine hamsters each were intranasally infected with two SARS-CoV-2 variants i.e., UK variant (hCoV-19/India/NIVP1 20203522/2020) and D614G variant (hCoV-19/India/2020770/2020) with 0.1 ml of virus of 10^5.5^ TCID50/ml under isoflurane anaesthesia and were used as donor hamsters for studying transmission via direct contact, aerosol and fomite routes ^5,6^. All the experiments were performed in triplicates for both the variants. SARS-CoV-2 genomic RNA (gRNA) load were tested in the nasal wash, throat swab and faeces of the contact hamsters on every alternate day till 14 days post exposure (DPE) using quantitative real-time RT-PCR for E gene as described earlier ^7^. Virus titration was also performed for the nasal wash samples in Vero CCL81 cells. The donor hamsters infected with UK variant showed progressive weight loss with the maximum average weight loss of 11 ± 2.82 % [mean ± standard deviation(S.D)] on day 6 whereas the donor hamsters infected with D614G variant showed a maximum average weight loss of −6.66± 1.36 % on day 8. The donor hamsters showed regain of weight thereafter. All the donor hamsters, bedding, cage surfaces and water bottle nozzle samples from cages used for fomite transmission study were tested for gRNA to ensure the presence of the virus before exposure (Table 1).

**Table 1:** SARS-CoV-2 viral gRNA load of donor hamsters and the cages used in the study at 24 hours post virus infection.

| Study | Sample details | Sample | Log<sub>10</sub> viral gRNA copies/ml |  |
| --- | --- | --- | --- | --- |
|  |  |  | VOC<br>202012/01 | D614G variant |
| Direct transmission | Donor hamster 1 | Throat swab | 8.40 | 7.83 |
|  |  | Nasal wash | 9.85 | 9.29 |
|  |  | Faeces | 8.33 | 7.10 |
|  | Donor hamster 2 | Throat swab | 7.66 | 7.90 |
|  |  | Nasal wash | 10.24 | 9.40 |
|  |  | Faeces | 7.51 | 4.99 |
|  | Donor hamster 3 | Throat swab | 8.01 | 7.39 |
|  |  | Nasal wash | 9.64 | 9.11 |
|  |  | Faeces | 5.10 | 6.22 |
| Aerosol transmission | Donor hamster 1 | Throat swab | 7.31 | 8.41 |
|  |  | Nasal wash | 9.68 | 10.03 |
|  |  | Faeces | 5.31 | 7.06 |
|  | Donor hamster 2 | Throat swab | 8.71 | 7.65 |
|  |  | Nasal wash | 10.40 | 9.86 |
|  |  | Faeces | 7.73 | 6.79 |
|  | Donor hamster 3 | Throat swab | 7.09 | 7.95 |
|  |  | Nasal wash | 9.84 | 9.38 |
|  |  | Faeces | 6.96 | 6.72 |
| Fomite transmission | Donor cage 1 | Bedding | 6.67 | 6.05 |
|  |  | Cage surfaces & water bottle nozzle | 7.24 | 6.91 |
|  | Donor cage 2 | Bedding | 6.36 | 5.00 |
|  |  | Cage surfaces & water bottle nozzle | 8.33 | 6.59 |
|  | Donor cage 3 | Bedding | 6.19 | 5.59 |
|  |  | Cage surfaces & water bottle nozzle | 7.13 | 6.73 |

Twenty four hour post infection, 3 donor hamsters were co-housed with a naive hamster (referred to as contact hamster further) in 1: 1 ratio in a new cage to study direct transmission and were observed till 14 DPE for body weight change and any respiratory signs. The contact hamsters exposed with UK variant showed maximum average weight loss of −2.93 ± 0.34 % and with D614G variant showed −4.8± 3.13 % on 8 DPE (Fig. 1A). The contact hamsters exposed with both variants showed viral gRNA positivity in the samples from 2^nd^ DPE and peak average viral gRNA load by 2^nd^ to 4^th^ DPE (Figure 1B-D). This is similar to the pattern of detection reported in intranasal inoculated hamsters with both variants ^3,8^. Titration of nasal wash samples showed consistent presence of virus till 10DPE with comparable titre in hamsters exposed with both variants (Fig. 1E).

**Figure 1:**
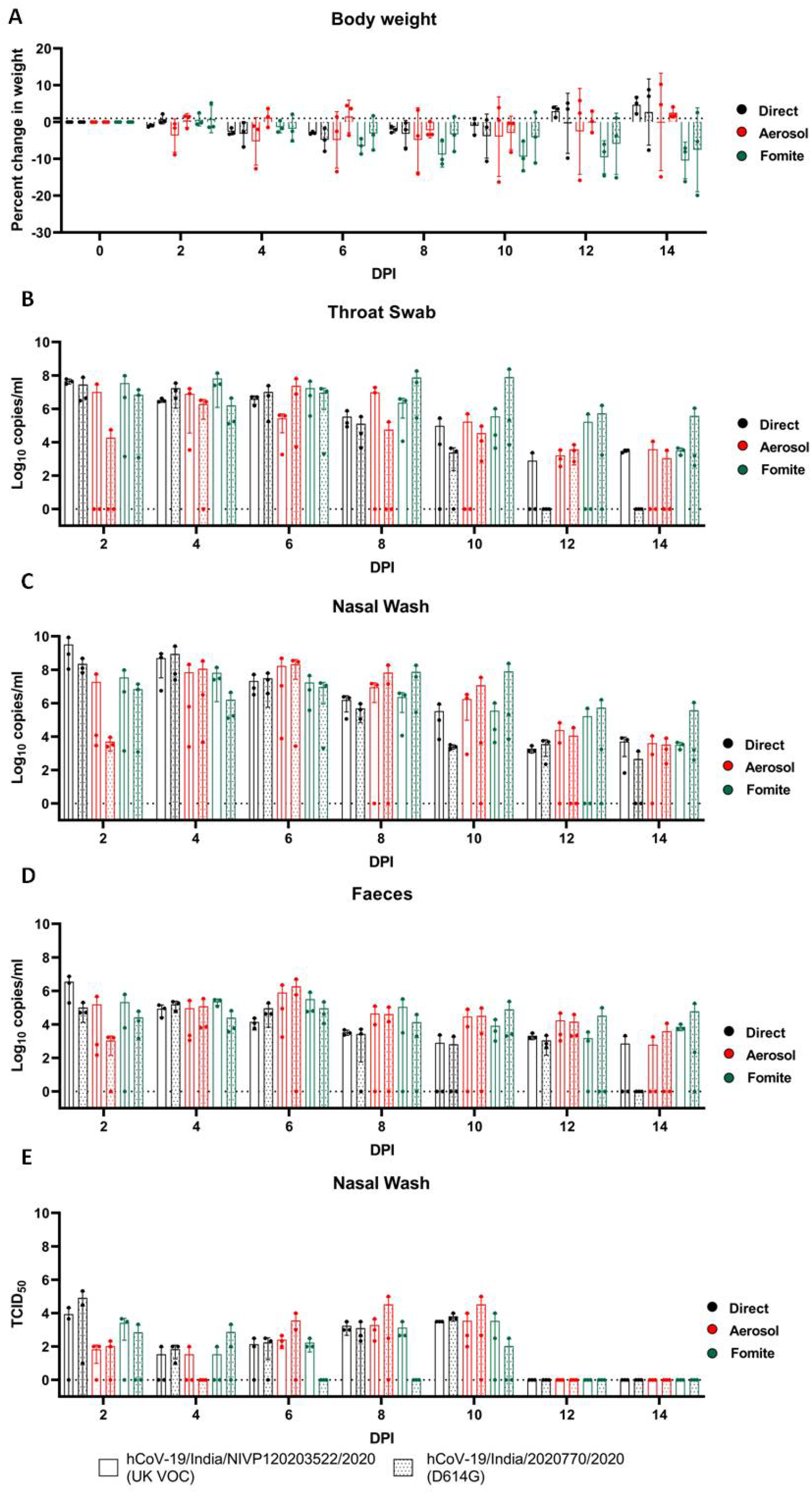
Percent body weight change and SARS-CoV-2 load in the hamsters post exposure. **(A)** The body weight change in hamsters following exposure with SARS-CoV-2 infected hamsters by direct, aerosol and fomite contact. Viral gRNA load in (B) throat swab (C) nasal wash (D) faeces in hamsters exposed by direct, aerosol and fomite contact. (E)Viral load in nasal wash samples of contact hamsters exposed by direct, aerosol and fomite contact estimated by titration in Vero CCL-81 cells expressed in TCID50.

As the direct contact transmission, could be contributed by aerosol and fomites, we assessed these routes of transmission alone. To assess the aerosol transmissibility, SARS-CoV-2 infected hamster after 24 hours post infection was co-housed in a modified individual ventilated cage with partition (which allows airflow) with a naive hamster for 8 hours. The contact hamsters were housed in new cages and were observed for 14 days. One hamster from the aerosol contact group of D614G variant did not show any body weight loss and showed very low or negligible amount of viral RNA load in the samples. The other hamsters of the aerosol contact group of UK variant and D614G variant showed a peak average weight loss of −8.9% on 8 DPE and −4.2% on 10 DPE respectively (Fig. 1A). The viral gRNA detection was observed from 2^nd^ DPE and the average viral gRNA peak detection in the nasal wash and faeces was observed on 6 DPE for hamsters exposed with both variants and higher viral load till 10 days (Fig. 1B-D). The TCID50/ml of the nasal wash samples from 6 to 10DPE showed higher titres (Fig.1E).

For the fomite transmission study, 3 naive hamsters were housed in different cages with soiled bedding of SARS-CoV-2 infected hamster housed for 48 hours following infection. A progressive decrease in body weight till 14 days was observed in fomite contact hamsters of both the variants (Fig. 1A). The peak average viral gRNA in fomite contact hamsters varied from 6 to 8 DPE in case of UK variant and 4 to 6 DPE with D614G variant (Fig 1B-D). This is in contrary to an earlier study which reported less efficient fomite transmission by SARS-CoV-2 in hamsters ^4^. Even though viral gRNA could be detected till 14DPE, live virus particles could not be detected from 6DPE in case of D614G variant contacts in contrast to UK variant contacts where it could be detected till 10DPE (Fig.1E).

The virus detection in contact hamsters were seen as early as on 2^nd^ DPE for both variants indicating faster transmission and a maximum infectious period of 10 days by all routes. The peak viral gRNA levels were comparable among different transmission routes whereas the peaks of detection by the aerosol and fomite route were found extended and with more variations among contact animals. The percent body weight loss also varied among different routes of transmission. This could be due to difference in the amount of virus dose exposure by these routes. Earlier research have shown that lower and higher dose of virus inoculums show comparable viral gRNA loads in hamsters but the lung lesions and body weight loss vary with virus dose ^3,9^.

IgG antibodies could be detected in all the donor and contact animals by an anti-SARS CoV-2 IgG ELISA on 21 DPE except one contact hamster of the D614G variant contact of the aerosol group which showed negligible viral gRNA load^10^. The day wise comparison of the body weight loss and the viral shedding pattern in contact hamsters by both variants on Mann-Whitney test did not show any statistical significance. Also the comparison of different routes of transmission by each variant on Kruskal Wallis test also did not show any statistical significance.

In conclusion, the transmission of SARS-CoV-2 variants could be established in Syrian hamsters by direct, aerosol and fomite routes as evident by the body weight loss, detection of viral gRNA/live virus in the samples and IgG antibodies in the contact hamsters. The study demonstrated comparable transmission efficiency of both UK and D614G variants of SARS-CoV-2 in Syrian hamsters.

## Acknowledgements

The authors acknowledge the support received from Prof (Dr.) Priya Abraham, Director, ICMR-NIV, Pune and the laboratory team of Maximum Containment Laboratory, ICMR-NIV, Pune which includes Mr Manoj Kadam, Mr. Annasaheb Suryawanshi, Mr Abhimanyu Kumar, Dr Rajlaxmi Jain, Mr. Rajen Lakra, Mr. Shreekant Baradkar, Mrs. Ashwini Waghmare, Ms. Manisha Dudhmal. Mr. Kundan Wakchuare, Ms. Tejashri Kore, Ms. Shilpa Ray, Ms. Priyanka Waghmare, Ms. Poonam Bodke. Authors also acknowledge Dr. SSYH Qadri, Head, Animal Facility and Dr. M. Satya Vani, ICMR-National Institute of Nutrition, Animal Facility for their support.

## References

1 WHO | SARS-CoV-2 Variants. WHO. http://www.who.int/csr/don/31-december-2020-sars-cov2-variants/en/ (accessed 8 Mar2021).

2 Davies NG, Barnard RC, Jarvis CI et al. Estimated transmissibility and severity of novel SARS-CoV-2 Variant of Concern 202012/01 in England. medRxiv 2020; : 2020.12.24.20248822.

3 Mohandas S, Yadav PD, Nyayanit D et al. Comparison of the pathogenicity and virus shedding of SARS CoV-2 VOC 202012/01 and D614G variant in hamster model. bioRxiv 2021; : 2021.02.25.432136.

4 Sia SF, Yan L-M, Chin AWH et al. Pathogenesis and transmission of SARS-CoV-2 in golden hamsters. Nature 2020; 583: 834–838.

5 Sarkale P, Patil S, Yadav PD et al. First isolation of SARS-CoV-2 from clinical samples in India. Indian Journal of Medical Research 2020; 151: 244.

6 Yadav PD, Nyayanit DA, Sahay RR et al. Isolation and characterization of VUI-202012/01, a SARS-CoV-2 variant in travellers from the United Kingdom to India. J Travel Med 2021. doi:10.1093/jtm/taab009.

7 Choudhary ML, Vipat V, Jadhav S et al. Development of in vitro transcribed RNA as positive control for laboratory diagnosis of SARS-CoV-2 in India. Indian Journal of Medical Research 2020; 151: 251.

8 Hou YJ, Chiba S, Halfmann P et al. SARS-CoV-2 D614G variant exhibits efficient replication ex vivo and transmission in vivo. Science 2020; 370: 1464–1468.

9 Ryan KA, Bewley KR, Fotheringham SA et al. Dose-dependent response to infection with SARS-CoV-2 in the ferret model and evidence of protective immunity. Nat Commun 2021; 12: 81.

10 Mohandas S, Yadav PD, Shete-Aich A et al. Immunogenicity and protective efficacy of BBV152, whole virion inactivated SARS-CoV-2 vaccine candidates in the Syrian hamster model. iScience 2021; 24: 102054.

